# Cultural signatures of punctuated environmental change: measuring the effects of population-wide stress on song distribution in dark-eyed juncos and song sparrows

**DOI:** 10.1101/2022.12.03.519005

**Authors:** Kate T. Snyder, Maria L. Sellers, Nicole Creanza

**Affiliations:** Department of Biological Sciences, Vanderbilt University, Nashville, TN; Evolutionary Studies Initiative, Vanderbilt University, Nashville, TN

## Abstract

Severe weather events can dramatically alter a species’ evolutionary trajectory. Previous landmark studies on populations in transiently hostile environments have measured traits that are under direct selection during those events. However, phenotype shifts that are not inherently adaptive to the fluctuating environment may also occur. Stress, especially during development, can cause important phenotypic changes in individuals, including impaired learning. Thus, learned behaviors, such as birdsong, may exhibit unique evolutionary dynamics as a result of widespread environmental stress. We hypothesize that ecosystem-level stressors may cause population-level changes to birdsong as entire cohorts experience developmental stress, learn songs imperfectly, and become the song tutors for future generations. In 2016, an unprecedented drought affected western New York State, a hotspot for community-science-generated birdsong recordings, and presented a unique opportunity to test our hypothesis through a natural experiment. We analyzed publicly available community-science birdsong recordings of two species, *Junco hyemalis* and *Melospiza melodia*, recorded between 2006-2020 in the drought-affected region and two control regions. We found that population-level song features had changed in the species with more complex songs (song sparrow) in the drought area after 2016, but not in the control area or in the species with a simple song (dark-eyed junco), implying that stress-induced deficits may disproportionately affect song traits that are more difficult to learn. These results demonstrate that environmental events can drive population-level trait evolution due to disruption in learning, with potentially important implications for speciation.

## Introduction

The natural world is punctuated by catastrophic events that can dramatically change natural selection pressures, altering the course of a species’ evolution [1,2]. The documented examples of these short-term phenomena on a population scale are characterized by unusually high mortality rates after all individuals are subjected to extremely hostile conditions, where those individuals that exhibit certain adaptive traits disproportionately survive to pass on their genes to the next generation [3–5]. These studies have centered on traits that are assumed to be genetically determined and not plastic in adulthood; however, recent research has shown that gene-by-environment interactions can have significant outcomes on phenotypes, which could theoretically also drive a species’ evolution, since natural selection primarily acts on expressed phenotypes. There are several potential mediators of phenotypic change based on interactions between organisms and their environments, including epigenetic modifications, microbiome composition, and learned or conditioned behaviors [6–8]. These factors can all be affected by the physical and social environment, especially those experienced during development, in potentially long-lasting ways that can be adaptive, conditionally adaptive in a given environment, or maladaptive [9–12].

Birdsong is a learned behavior that is extremely diverse across the 4000+ species that exhibit it in the songbirds (suborder: Oscine, Order: Passeriformes). Learning duration varies between species, with some species learning only in the first year of life, after which they do not modify their songs (termed “closed-ended learners”), and others learning for several years or throughout their lifetimes (“open-ended learners”). Song evolution is influenced by sexual selection, as it primarily functions in mate attraction, species identification, and territory defense. Songs are composed of syllables, defined as periods of continuous sound that are separated by a short period of silence. The diversity of song is typically characterized using a set of metrics that include measurements of pitch, frequency modulation, duration, spectral entropy, and syllable diversity. One key metric is “syllable repertoire,” or the total number of unique syllables an individual produces. The average syllable repertoire within a species ranges from 1 (e.g. in the chipping sparrow [13]) to over 2000 (in the brown thrasher [14]), and is considered a metric of song complexity and learning capacity [15,16]. The song preferences of females vary widely between species, but in laboratory and field studies across several species, it has been found that females tend to prefer more complex songs, as measured by a male’s syllable repertoire, number of unique syllables per song, or number of unique songs [17–19].

Birdsong is an energetically costly behavior, requiring significant investment in specialized neural development and time dedicated towards practicing song [20]. Stress during development has been shown to impair song learning ability in several species, using song learning accuracy or song complexity as proxy metrics [21,22]. The developmental stress hypothesis posits that birdsong is an honest indicator of fitness because females can gain accurate information about a male’s developmental environment by assessing song quality, enabling them to choose a mate that had a non-stressful upbringing [20]. The conditions causing stress, such as food availability and parasite load, are usually assumed to vary across a population, with some birds experiencing little stress during development, and attractive song characteristics are favorably selected by females. However, certain extreme environmental events might effectively apply stress to an entire population simultaneously, which could result in an entire generation of birds with impaired song learning.

Genetically determined phenotypes in a population are constantly undergoing minor changes as individuals exhibit varied success from year to year, but major shifts in population phenotype happen far more rapidly under extreme conditions [1]. Could a similar phenomenon occur in a culturally transmitted phenotype such as song that does not give a selective advantage under the environmental change? Specifically, if a stressor was applied across an entire breeding songbird population simultaneously, such that virtually all juveniles experienced impaired learning, would there be a detectable perturbation in population song in the next breeding season? If these juveniles’ songs became the models for future generations, could such a punctuated stressor ultimately induce a lasting change in the populations’ songs? We predict that ecosystem-level stressors may cause population-level changes to bird song as entire cohorts that experience developmental stress subsequently learn their songs imperfectly. Further, if these individuals become the song tutors for future generations, these altered songs could theoretically spread in the population and become an established dialect.

While the biological effect of developmental stress upon the song quality of individuals has been well studied, the cultural effect of an environmental stressor applied to an entire natural population has not, to our knowledge, been investigated. We have identified an ecological opportunity through which to attempt to observe the effects of such a phenomenon. From June through November 2016, New York State, which typically has a humid climate, experienced a severe drought. The most severely affected region, central-western New York State, including Ithaca and the surrounding area, spent over two months under the classification of “Extreme Drought” [Information from the Drought Monitor [23], accessed via https://droughtmonitor.unl.edu/NADM/Home.aspx]. Air temperatures in 2016 in this region were also higher than normal, around the 90th percentile relative to the previous 60 years [24]. These combined factors had significant effects on vegetation, using agriculture losses as a proxy metric; in the most heavily impacted region, New York Drought Region VII, which encompasses Tompkins, Cayuga, Onondaga, Schuyler, Seneca, Yates, Ontario, Livingston, and Wyoming counties, farms reported average non-irrigated crop losses ranging from 45% loss of field crops to 69% loss of fruit crops [24]. Incidentally, this area is also a hotspot for songbird recordings: the area surrounding Ithaca, New York, which is home to Cornell University, the Cornell Lab of Ornithology, and the Macaulay Library of Natural Sounds, has a high density of publicly available bird song recordings obtained by both academics and community bird enthusiasts over the past few decades, with a sharp increase in recordings in the past five to ten years.

While there does not seem to be robust analysis or systematic documentation of effects on passerines during this drought, research on passerine species from other drought events can provide some predictions as to the potential effects. In many species, including song sparrows, reproduction and reproductive physiology is negatively affected by the absence of water [25–27]. This extends to decreases in nesting behavior and egg-laying [28]. Drought leads to physiological stress and mortality in individuals by reducing the availability of plant and arthropod biomass that serve as food sources for many bird species, as well as through dehydration [28,29]. These effects may be most pronounced in specialist species, but generalists are also affected negatively by lack of water availability and tree and plant mortality [30].

Here, we present analyses of the songs of two focal species, the dark-eyed junco (*Junco hyemalis*) and the song sparrow (*Melospiza melodia*), recorded in central-western New York State between 2006 and 2019. We analyze these recordings obtained from publicly accessible repositories with a specific prediction that the population distributions of certain song features will shift from before the 2016 drought to after the drought. We hypothesize in particular that metrics of song complexity in song sparrows will shift towards simpler song, since song complexity has previously been shown to negatively correlate with metrics of stress in song sparrows [31]. We predict that there will also be a larger variance in these feature distributions after the drought, since recordings after 2016 are likely to represent a mix of individuals that learned their songs prior to 2016, individuals that learned their songs during drought period, and individuals that may have learned their songs from the drought-year birds.

## Methods

### Site Selection

In the summer of 2016, a large area of New York State experienced a drought that was unprecedented in recorded history for the region. This provided a unique opportunity for a natural experiment to study the effects of environmental stress on birdsong on a population-wide scale. Tompkins County, home to the Cornell Lab of Ornithology, was classified as being under drought conditions from late June until well into November of that year, with over two months spent under the classification of “Extreme Drought.” Using the ample community-science birdsong recording data available in this area, collected before, during and after the drought, we quantify features of the songs of individual species to assess whether the drought had a detectable effect on songs in the affected populations. We compare the songs from the drought area with songs in a control region of approximately similar land area. The control regions were selected by finding the area with the next highest density of recordings after western New York from the pool of available recordings on Macaulay Library and xeno-canto. We assessed the density of recordings for each target species and both time periods, 2006-June 2016 (before drought) and 2017-2019 (after drought) and selected the regions that were not impacted or were less severely impacted by the 2016 drought and that had the greatest number of available recordings before and after the drought period. For both species, we only considered regions in the eastern half of the continental U.S. since there are well-documented differences in song dialects and singing behavior between east and west coast populations of song sparrows [32], and different dark-eyed junco subspecies are present in the West [33]. Since no regions outside of western New York had at least 20 recordings for dark-eyed juncos from 2006-2016, we chose the dark-eyed junco control region based on the abundance of recordings from 2017-2019, and, of those options, chose the region that did not also experience severe drought during 2016. (**Figures S1A-B**) For song sparrows, only one potential control area had relatively dense recording availability both before and after 2016, and it had experienced moderate drought during 2016. (**Figures S1C-D**)

**Figure 1.**
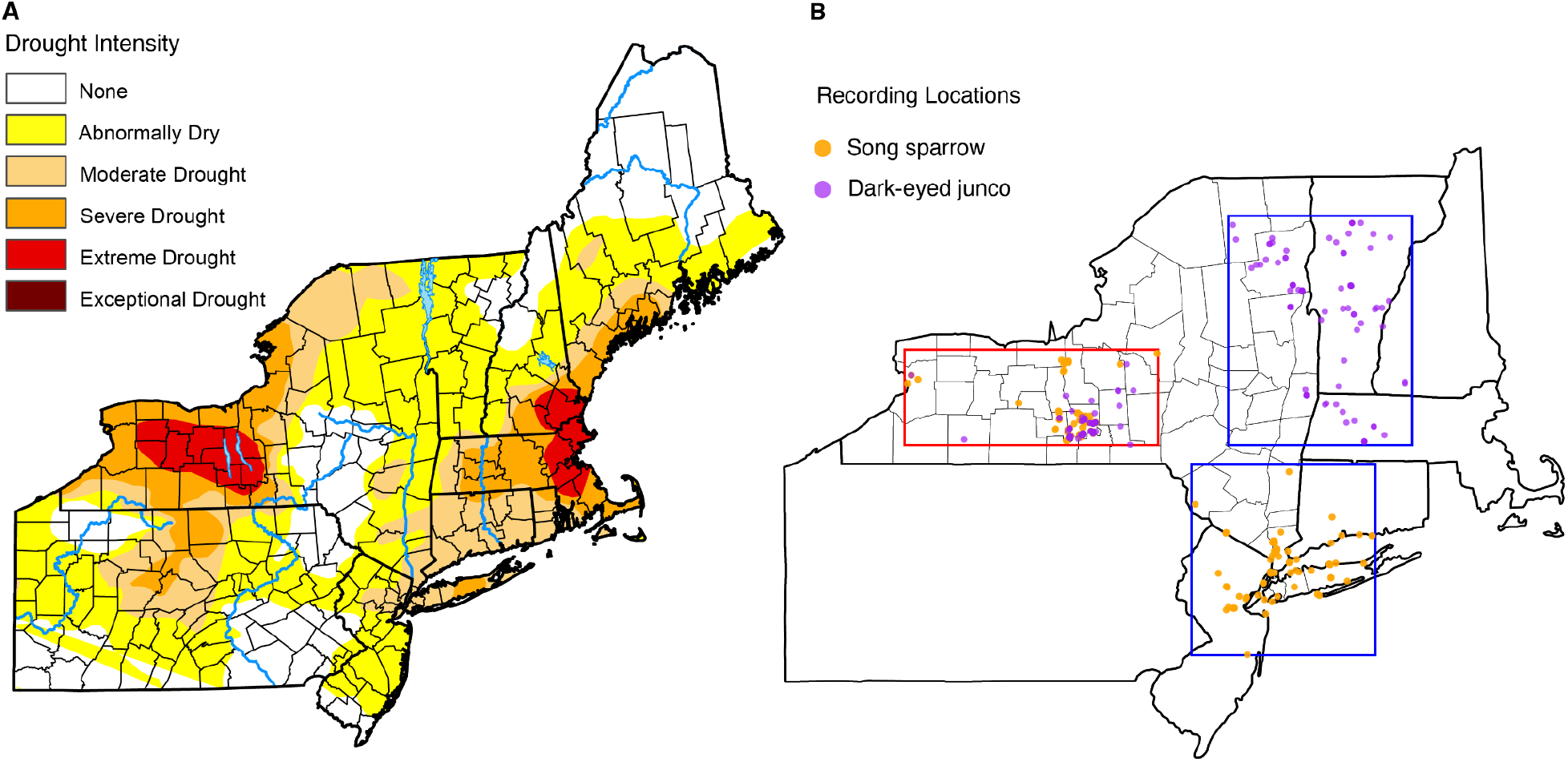
Maps of drought severity and recording locations. (A) Map from U.S. Drought Monitor [23] from September 2016 showing the severity and geographic extent of the drought in western New York. (B) Map of songs for song sparrows and dark-eyed juncos in our analyses. The red box encompasses the recordings of both song sparrows (orange) and dark-eyed juncos (purple) in the drought region. For a similarly sized control region that did not experience a drought, we found the densest sampling of song sparrow recordings in northern New Jersey and southern New York and Connecticut and of dark-eyed junco recordings in eastern New York, Vermont, New Hampshire, and western Massachusetts.

### Species Selection Criteria

We selected song sparrows and dark-eyed juncos as target species based on several factors related to 1) feasibility of study, 2) likelihood of drought impact on individuals, and 3) likelihood that any drought-associated impact of song could be detected in years following the drought. Regarding the feasibility of study, we considered the number of song recordings available and song structure. In early 2017, we retrieved metadata for all audio recordings from Tompkins County, New York which were tagged as containing “song” in the Macaulay Library online repository. We excluded any species with less than 100 recordings. We also eliminated any species that sing continuously, or that do not have a clear separation between song bouts to facilitate consistent song analysis using our acoustic analysis program, Chipper [34]. Drought-associated impacts on song within a population would be diluted by high rates of immigration or emigration, so we sought species for which there is limited dispersal or at least some evidence of philopatry, so that adult birds recorded in a location are likely to have hatched near that location. We also focused on species that do not modify their song repertoires after their first breeding season (i.e. “closed-ended” learners), since we hypothesized that effects of the drought on song learning could be less pronounced in “open-ended” learners, who can continue to modify their song in subsequent breeding seasons [35]. Based on these criteria, we selected the dark-eyed junco (*Junco hyemalis*) and the song sparrow (*Melospiza melodia*), the two songbird species with the most recordings in the drought area, both of which are closed-ended learners and show evidence of philopatry [36,37], suggesting that there would be continuity in population makeup in the area from before to after the drought.

Environmental limitations to food sources may have been a major contributor to developmental stress during the drought [38,39]. The 2016 drought has been shown to have had a large detrimental effect on agricultural vegetation and wild vegetation alike [24]. The diet of *Junco hyemalis* is composed primarily of plant matter, which comprises 76% of their diet, 62% of which is seeds, and they have been known to supplement their diet with insects on occasion [40]. Although the diet of *Melospiza melodia* is also primarily composed of plant matter for most of the year, during the breeding season their diets rely more heavily on animal food sources, such as beetles, butterflies, and small gastropods [40].

The two focal species for our study have notable variation in song complexity. The dark-eyed junco has a relatively simple and consistent song, typically consisting of one syllable sung repetitively, whereas the song sparrow sings many songs composed of multiple syllables. Dark-eyed juncos have small repertoires of around 4 songs on average, and have been known to sing the same song up to 120 times in a row before switching to a new song [41–43]. Song sparrows, on the other hand, typically cycle between 5-13 different song types, often singing several renditions of one type before switching to the next [44].

### Obtaining Recordings

Two citizen-science recording libraries, Macaulay Library and Xeno-canto, provided most of the recordings for this study. We utilized the search engines of these digital databases with the search terms “*Junco hyemalis*” and *“Melospiza melodia*” with the filters “Location = New York, United States’’ and “Sounds = Song.” From these digital databases, we were able to obtain a total of 203 recordings for dark-eyed juncos, and 564 recordings for song sparrows from within the two analysis regions from 2006-2019. After examining each recording, we discarded any that misidentified the species, only contained calls, were noted in the recording metadata to contain juvenile or plastic song, or had excessive background noise. In cases where multiple recordings existed from the same recordist on the same date and at the same time or location, we used only one recording in order to avoid double-counting individual birds who may have been recorded more than once, unless it was clearly noted in the remarks that the recordings were of different individuals.

We supplemented these recordings with ones recorded in the field by one author (KTS) in July 2017. During this trip, KTS visited several publicly accessible locations in Tompkins County, within the region that had experienced the heaviest drought during 2016. This added an additional 8 dark-eyed junco and 7 song sparrow recordings that were usable. Recordings from before 2006 through June 2016 were categorized as before the drought. Anything recorded from July 2016 through December 2016 was considered to have occurred during the drought, and thus removed from analysis. Any recordings from 2017 or later were defined as after the drought. Due to the sparseness of recordings in public repositories from earlier years, we did not include any recordings prior to 2006.

### Processing Song Recordings

Many of the recordings collected contained multiple song bouts from the same bird. For the purposes of this study, we define a bout as a continuous period of syllable production within the recording that is visibly separate from other periods of song. Within these bouts, syllables are separated from one another by silence, and discrete pulses of signal that are not separated by silence are termed notes (**Figure 2**).

**Figure 2:**
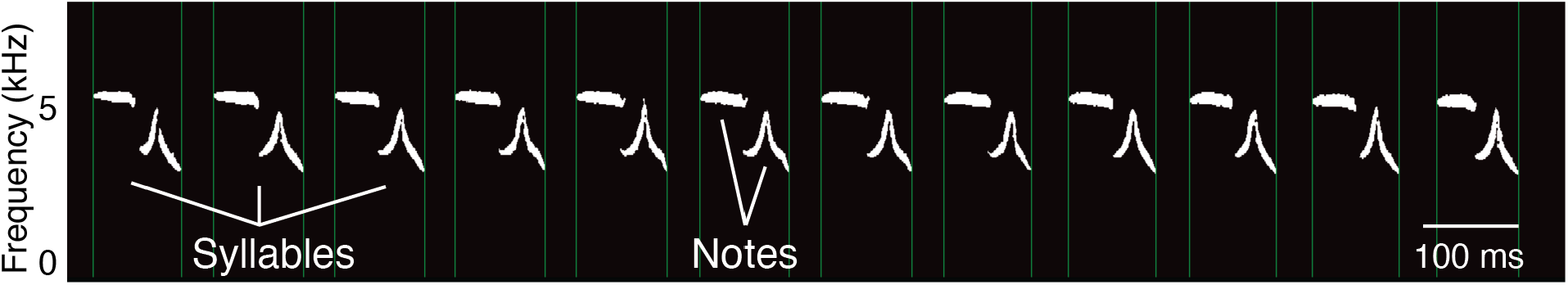
Example spectrogram of a single bout of dark-eyed junco song with parts of the song highlighted. Syllables are periods of signal separated by silences. Within these syllables, notes are the discrete pulses of signal that are not separated from one another by silences.

Individual bouts were extracted from recordings using Audacity version 2.4.2 and exported as separate WAV files with sampling rate set to 44100 Hz. Due to inherent differences between species song, different bout-selection protocols were implemented for juncos and song sparrows. Since dark-eyed juncos repeat the same song type many times in a row, most of the recordings in community-science databases are of a single song type [41,42]. If there were multiple distinct song types in a single recording, we collected at least one bout of each type, but otherwise, we collected up to three bouts per recording, preferentially choosing ones that were least affected by background noise. Song sparrows, in contrast, cycle between song types more frequently [44]. In order to obtain the greatest possible variation in songs, we parsed out every available bout from song sparrow recordings.

### Syllable Segmentation in Chipper

Bouts were segmented into discrete syllables by a researcher blind to the region and year of each recording using the open source software Chipper [34]. Chipper uses a Fast Fourier Transform to generate a spectrogram from the WAV file of each song bout and then predicts syllable boundaries using fluctuations in the signal amplitude. The user can first implement high-pass and low-pass filters to reduce background noise and then modify the predicted syllable boundaries by adjusting parameters such as the signal-to-noise threshold, the minimum silence duration required to distinguish between syllables, and the minimum duration of syllables. Song bouts were discarded if they had a signal-to-noise ratio that made it difficult to reliably detect syllable boundaries. In addition, any files that had birds singing in the background that interfered with the song of the bird of interest in the recording were removed. Since dark-eyed juncos generally sing a repeated syllable, we could extract syllable properties even when only a subset of the song bout was usable; these files were excluded from analyses of two song features, number of syllables and bout duration.

The resulting spectrogram of each song is stored as a matrix in which (at a sampling rate of 44100 Hz) every row represents ∼43 Hz and every column represents 0.317 milliseconds [34]. The value in each element of this matrix represents the signal intensity at that frequency and time; each element in the matrix becomes a pixel in the spectrogram image (e.g. **Figure 2**). After the user adjusts the signal-to-noise threshold, signal intensities above this value are retained in a binary spectrogram where ‘1’ or ‘0’ indicates that signal is present or absent at that frequency and time.

Chipper also executes an additional stage of noise reduction by measuring each note, defined as an area of continuous sound across the time and frequency axes, and discarding any that have an area in pixels less than a user-specified noise threshold. Chipper allows the user to empirically determine this average noise threshold (the minimum area in pixels that a note must be in order to not be discarded as noise) as well as an average syllable similarity threshold (the percent overlap that determines whether two syllables are considered the same or different). To assess these thresholds, we selected ∼80-100 single-bout files for each species and manually set the noise threshold and syllable similarity threshold for each file. Across these test files, the average noise threshold for song sparrows was 59.1 pixels, and the average similarity threshold was 47.3% syllable overlap. The average noise threshold for dark-eyed juncos was 142 pixels, and the average similarity threshold was 29% syllable overlap. We used these average values as the thresholds in subsequent analyses. Occasionally, a syllable would only contain notes with areas below the noise threshold; we manually checked these recordings to decide whether the songs should be re-analyzed with different parameters or discarded.

### Song analysis

After syllable segmentation, we used Chipper to measure a slightly different set of song features for dark-eyed juncos and song sparrows, choosing metrics that were more likely to be meaningful based on the structure and features of each species’ song. For both species, we measured bout duration, total number of syllables, number of unique syllables, total number of notes, number of notes per syllable, mean syllable duration, standard deviation of syllable duration, rate of syllable production (calculated as the number of syllables divided by the duration of the song bout), degree of syllable repetition (calculated as the number of syllables divided by the number of unique syllables), overall frequency range, average maximum frequency of syllables, and average minimum frequency of syllables. For dark-eyed juncos, which generally sing multiple repetitions of the same syllable, we also measured the mean syllable stereotypy, defined as the mean syllable overlap value for syllables that were deemed to be repetitions of the same syllable. For song sparrows, which generally sing both short and long syllables, we also measured the duration of the shortest syllable and the duration of the longest syllable. Features that describe duration are measured in milliseconds, and features that describe frequency are measured in Hertz.

For each focal species and song feature, we used a set of Shapiro-Wilk tests to assess the distribution of our song-feature data before and after the drought in both the drought region and the control region. When at least one group in each comparison was not normally distributed (**Table S2**), we log-transformed the song-feature data. If the log-transformed values for that song feature were still not normally distributed, we analyzed the song features using non-parametric statistical tests, i.e. using a Wilcoxon rank-sum test instead of a *t*-test to compare data before versus after the drought in both the drought region and the control region, with a Holm-Bonferroni correction for multiple hypotheses. We also assessed the homogeneity of variance before versus after the drought in both regions using Brown-Forsythe tests. All statistical analyses were implemented in R with custom code, available at github.com/CreanzaLab/Birdsong_DroughtStress.

## Results

### Dark-eyed junco song analysis

We analyzed 403 dark-eyed junco bouts from 175 unique recordings in total (Table S1). We discarded 28 of our original 203 recordings; these were either recorded during the drought (5 recordings), recorded slightly outside the drought region, or had no analyzable song due to excessive noise or poor signal-to-noise ratio. In the drought region, we analyzed 46 recordings (131 bouts) from before the 2016 drought and 73 recordings (199 bouts) from after. From the control region, where the drought was less severe or absent, we analyzed 8 recordings (11 bouts) from before 2016 and 48 recordings (62 bouts) from after 2016.

Using Chipper, we segmented these dark-eyed junco song bouts into syllables and measured thirteen properties from each: bout duration, number of syllables, number of unique syllables, total number of notes, degree of syllable repetition (equal to the total number of syllables divided by the number of unique syllables per bout), mean number of notes per syllable, rate of syllable production, mean syllable duration, standard deviation of syllable duration, mean syllable stereotypy, frequency range, average maximum frequency of syllables, and average minimum frequency of syllables.

If a recording had multiple bouts, we took the mean value of each song feature across bouts so that each recording would be weighted equally in the comparison. For both the drought region and the control region, no song features were significantly different before vs. after the 2016 drought (**Figure 3, Table S2**). In addition, we did not reject the null hypothesis that each song feature had equal variance before versus after the drought (**Table S2**).

**Figure 3.**
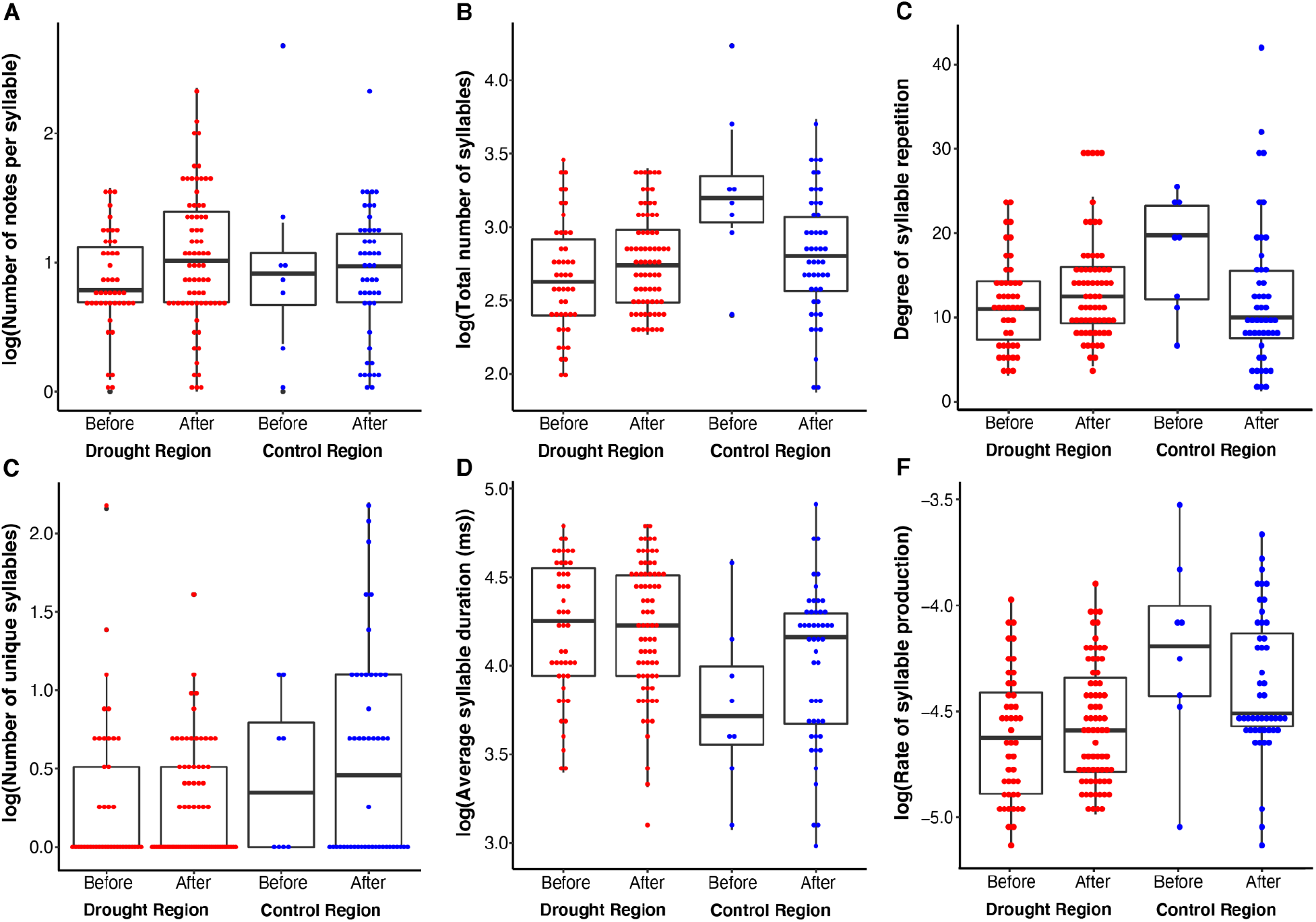
Song feature distributions within dark-eyed junco populations before and after the 2016 drought. Each boxplot shows dark-eyed junco mean song-feature values per recording: (A) number of notes per syllable, (B) number of syllables, (C) degree of repetition (number of syllables divided by the number of unique syllables), (D) number of unique syllables, (E) mean syllable duration, and (F) rate of syllable production.No dark-eyed junco song feature significantly differed in median or variance when we compared recordings before versus after the drought. Note that we log-transformed song features when they were not normally distributed (all panels except C). The midline of each boxplot represents the median, with the boxes representing the interquartile range (IQR). The whiskers are 1.5 * IQR. These specifications also apply to the box plots in Figure 6.

The dark-eyed junco sings a relatively simple song characterized by a single syllable repeated multiple times. We sought to visualize a ‘typical’ song from each of the four groups: before and after the drought from the drought region and before and after the drought from the control region. We identified the most representative bout for each group by first rescaling the song features with the largest differences from before to after the drought for each bout—the number of notes per bout, bout duration, and degree of repetition—so that their values would be comparable to one another (base R function ‘scale’). We calculated the average value for each scaled feature among the bouts in each recording, and found the median value of each feature across recordings within each group. (**Figure 4**). We then identified and plotted the spectrograms of the bouts that were the smallest Euclidean distances from the point located at the median value of the three song features for each group (**Figure 5**).

**Figure 4.**
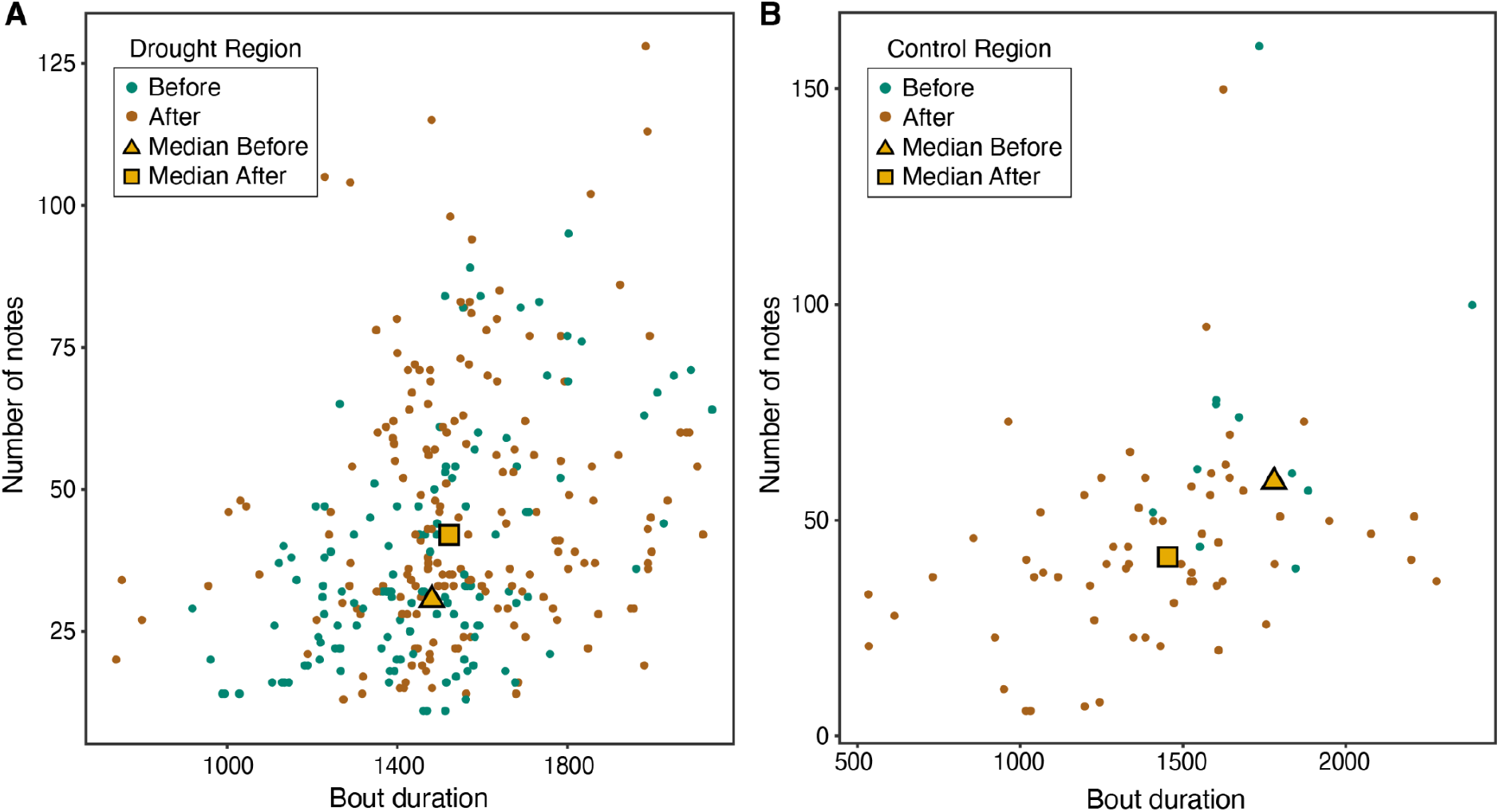
Visualizing dark-eyed junco song features. For every analyzed dark-eyed junco bout, we visualized the two song features with the largest differences before versus after the drought, number of notes and bout duration, in both A) the drought region and B) the control region (**Table S2**), though these differences did not reach statistical significance with a Holm-Bonferroni correction. The yellow shapes indicate the median value of both of these song features.

**Figure 5.**
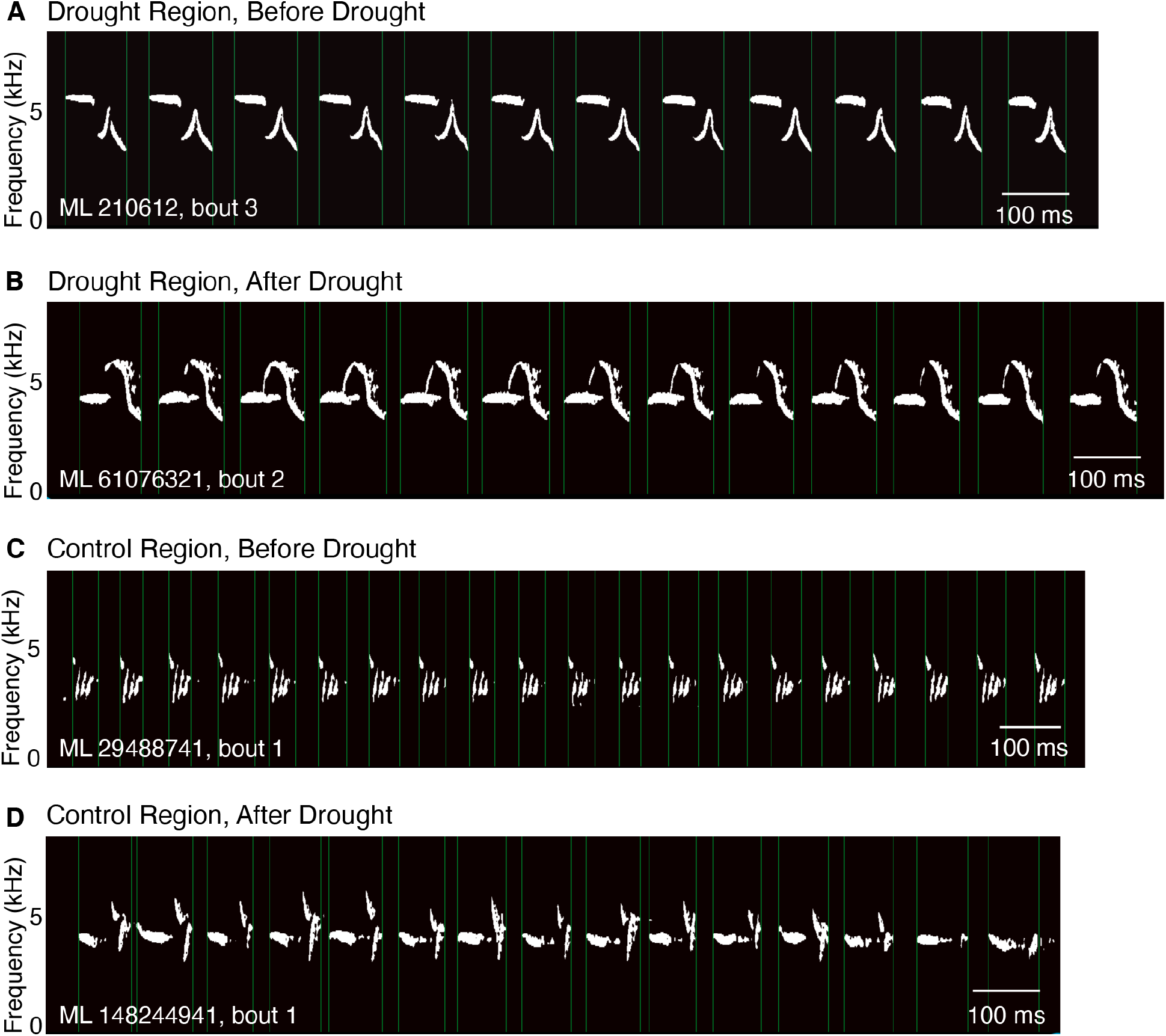
Spectrograms from representative dark-eyed junco songs. The songs that were closest to the median values of the three song features that showed the greatest difference before versus after the drought—the total number of notes, bout duration, and degree of repetition—are shown for each region. The groups that these songs represent did not significantly differ from one another in any of the measured song features.

### Song sparrow song analysis

We analyzed 1556 song sparrow bouts from 295 unique recordings in total (**Table S3**). We discarded 269 of our original 564 recordings; numerous recordings were submitted by the same recordist (or a group of recordists in the same location) on the same day, and we eliminated all but one from our sample unless we could confirm that the recordings were different birds based on the information provided by the recordists. In addition, we eliminated songs that were either recorded during the drought (7 recordings), recorded slightly outside the drought region, or had no analyzable song due to excessive background noise or poor signal-to-noise ratio. In the drought region, we analyzed 37 recordings (409 bouts) from before the 2016 drought and 159 recordings (745 bouts) from after. From the control region, where drought conditions reached only moderate severity, we analyzed 16 recordings (72 bouts) from before 2016 and 83 recordings (330 bouts) from after 2016.

For each of the fifteen song features we analyzed for song sparrows, we calculated the mean value across bouts in each recording and compared recordings before versus after the 2016 drought in both the drought and control regions. In the drought region only, three song features were significantly different after the 2016 drought: the number of syllables (*p* < 2.6×10^−3^), the number of notes per syllable (*p* < 5.9×10^−4^), and the degree of repetition (*p* < 5.0×10^−5^) (**Figure 6, Table S3**). However, as with the dark-eyed junco, we did not reject the null hypothesis that each song feature had equal variance before versus after the drought (**Table S3**).

**Figure 6:**
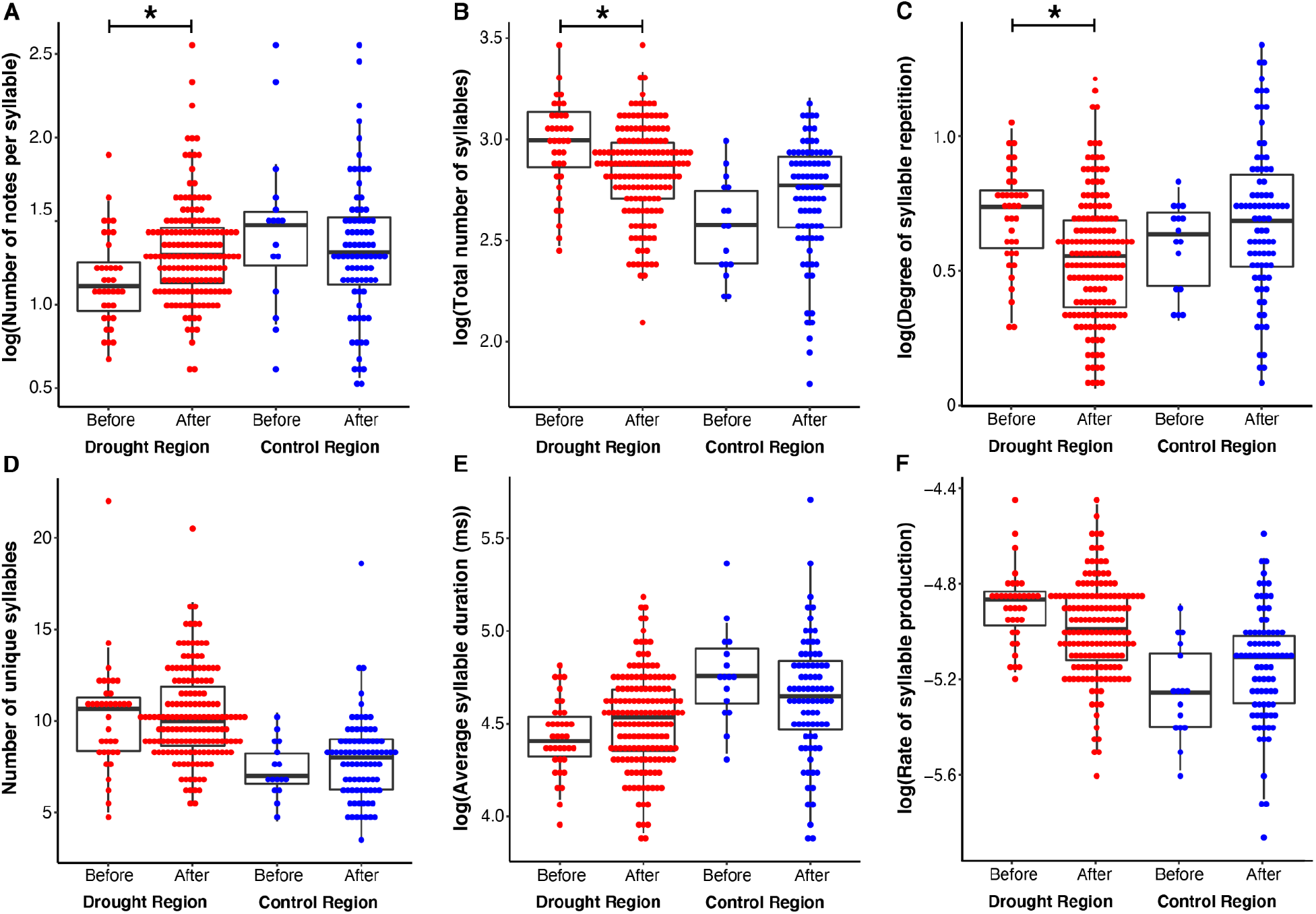
Song feature distributions within song sparrow populations before and after the 2016 drought. Each boxplot shows song sparrow mean song-feature values per recording: (A) number of notes per syllable, (B) number of syllables, (C) degree of repetition (number of syllables divided by the number of unique syllables), (D) number of unique syllables, (E) mean syllable duration, and (F) rate of syllable production. The number of notes per syllable, number of syllables, and degree of repetition (A-C) were significantly different from before to after the drought in the drought region only, with Bonferroni-Holm correction for multiple hypothesis testing. In the drought region, the difference in the rate of syllable production from before to after the drought had a Wilcoxon rank-sum test *p* = 0.0074, but was not significant after Bonferroni-Holm correction for multiple hypothesis testing. The number of unique syllables was normally distributed (Shapiro-Wilk test *p* > 0.05) in both time periods and did not need to be log-transformed. (**Table S3**)

Song sparrows sing a relatively complex song, particularly for closed-ended learners [45]. Many of their syllables are composed of multiple notes, and they cycle through their syllable repertoire of approximately 35-38 syllables by singing different song types containing overlapping subsets of their syllable repertoire. To capture this variation, we analyzed as many bouts as possible from each individual recording. Bout-level data from the two song features that showed the strongest temporal difference in the drought region, the number of notes per syllable and the number of syllables per number of unique syllables, are shown in **Figure 7**. To visualize a ‘typical’ song bout from each region and each time period, we rescaled the three significantly different song features so that they would be comparable with one another, and we found the song bout that was closest to the median value of all three features (**Figure 8**).

**Figure 7.**
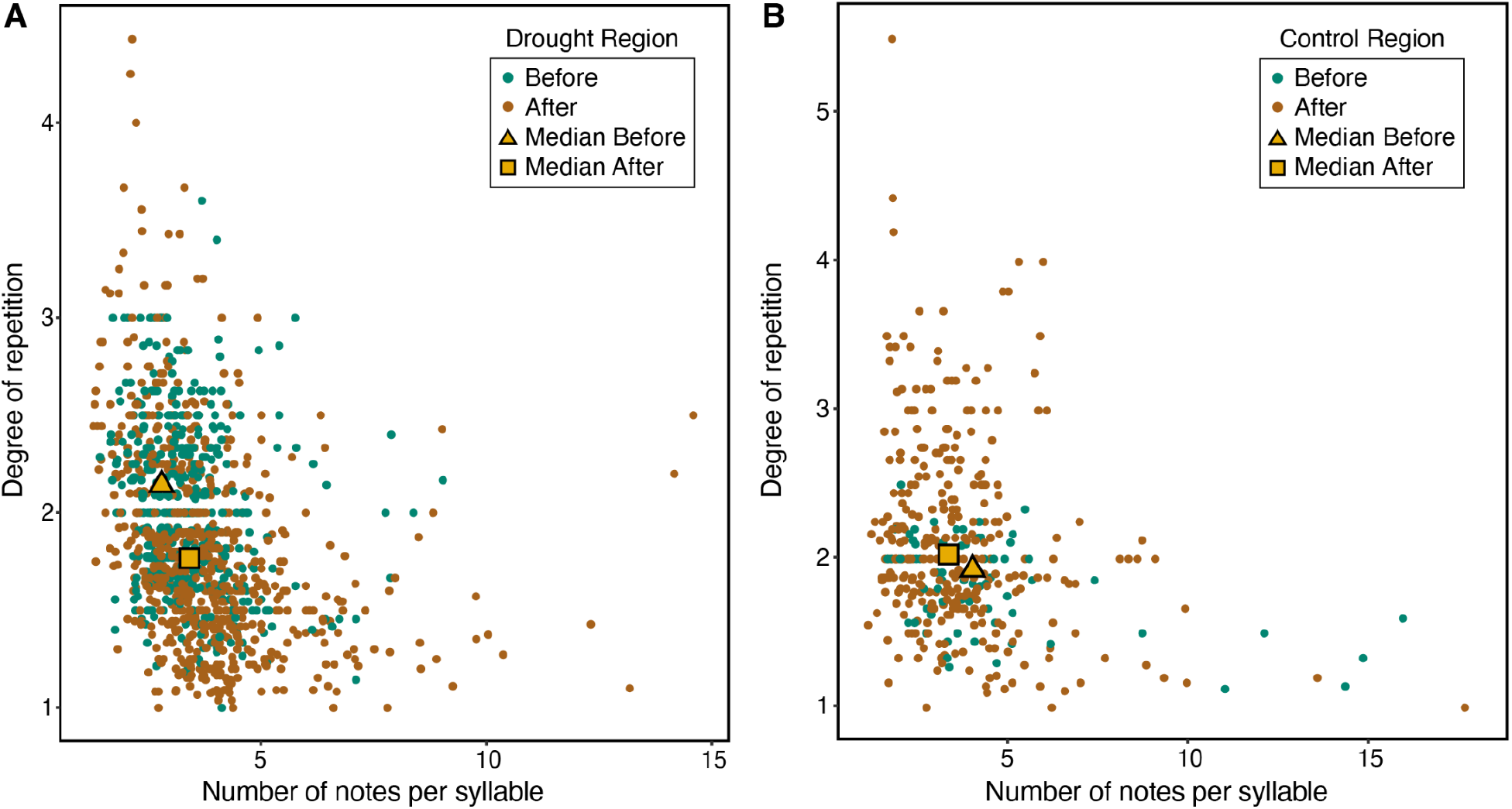
Visualizing song sparrow song features. For every analyzed song sparrow bout, we visualized the two song features with the largest differences before versus after the drought, number of notes per syllable and degree of repetition (**Table S3**), both of which were significantly different in the two time periods in the drought region (A) but not in the control region (B). After the drought, song sparrow songs on average had less syllable repetition and more notes per syllable. There were no significant differences in the control region.

**Figure 8.**
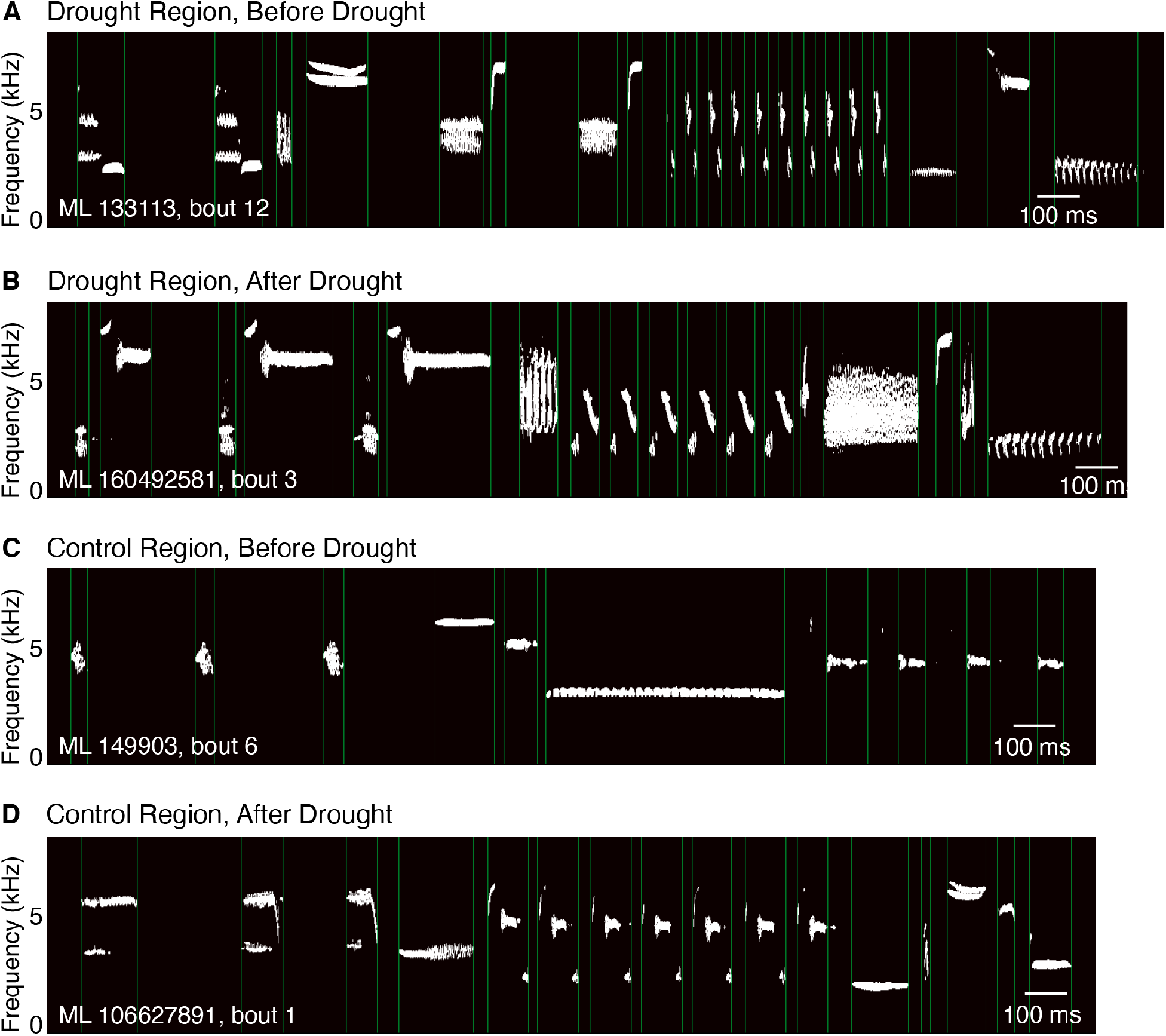
Spectrograms from representative song sparrow songs. These songs were closest to the median rescaled values of the three song features that showed significant differences before versus after the drought in the drought region, as in **Figure 7**. Compared to before the drought, song sparrow songs after the drought had significantly more notes per syllable, fewer total syllables per bout, and a lower degree of repetition. Here, the ‘median’ song from before the drought has 21 syllables and 10 unique syllables, with a degree of syllable repetition of 2.1; the ‘median’ song after the drought has 18 syllables and 10 unique syllables, thus a degree of syllable repetition of 1.8.

## Discussion

Here, we analyze community-science birdsong recordings from two common species of songbird, the dark-eyed junco and the song sparrow. In particular, we compare the time period before and after a severe drought in western New York, and compare these results to songs from control regions, in which drought was absent or only moderate, over the same time period (**Figure 1**). Because physiologically stressful conditions during development can disrupt song-learning on an individual level, we hypothesized that a population-wide stressor could have measurable effects on the song distribution of a population. For dark-eyed juncos, a species that generally sings a simple song of one repeated syllable, we observed no significant changes in any measured song feature in either the drought region or the control region. In the song sparrow, a species that has a relatively large syllable repertoire and produces complex multi-note syllables, we observed significant differences in song features related to both syllable complexity (notes per syllable) and song complexity (number of syllables, degree of repetition) in the drought region, but not the control region. Specifically, song sparrows after the drought sang songs with more notes per syllable, less repetition, and fewer syllables overall. It is notable that we see these significant effects in the species with more complex songs, since we hypothesized that species with complex songs would be more sensitive to the potential stress-associated learning disruption. However, the behavioral consequences of the changes we observed are not completely clear; more notes per syllable after the drought is suggestive that syllables in the later group might be more complex, whereas fewer overall syllables per bout could suggest reduced song complexity. The reduction in the degree of repetition after the drought (**Figure 6C**) is less straightforward, but when taken in concert with the trends towards longer average syllable durations and a lower rate of syllable production (**Figures 6E, F**), it may indicate that the shift occurs most prominently in the trill section that exists in most song sparrow songs, in which a single syllable is rapidly repeated on average 7-9 times [46]: a smaller number of syllable repetitions in the trill would decrease the overall degree of repetition and, if the trill was approximately the same duration, the rate of syllable production in the trill. We performed a follow-up analysis to check whether this interpretation is borne out in the data by obtaining for each bout the number of times the most-repeated syllable type was produced in that bout. We averaged these values across all bouts within each recording to obtain, for each recording, the “mean number of repetitions of the most-repeated syllable type per bout” and found that this was significantly lower after the drought in the drought region only **(Figure S2)**. This provides further evidence that trills may have meaningfully changed due to the drought, becoming either shorter in duration or slower in the rate of syllable production. Notably, a lower rate of syllable production could be indicative of poorer performance: though the characteristics that determine trill quality haven’t been studied intraspecifically in song sparrows, trills, especially ones that cover large frequency ranges, are considered to be physically difficult to perform across species, and thus a higher rate of trill production is considered to be indicative of higher quality or greater skill [47]. There is evidence from several other species that trill rate is important in female choice, territory defense, and intrasexual conflict [48–51]. Further, there is some evidence that rate of syllable production may be affected by developmental history in swamp sparrows, a close relative of song sparrows [51].

Song sparrows have a long study history in the field of songbird neuroethology, providing additional context for these observations. There have been several lab-based song-preference studies in wild-caught song sparrows that have found that females exhibit stronger preferences for familiar song types than unfamiliar types [52–54] and for larger song repertoires [18]. In field studies, males with larger song repertoires held territories longer and experienced greater reproductive success [55], as well as had a higher probability of mating in their first breeding year [19], although other studies have not found such relationships between song repertoire and reproductive success in the field [18]. Studies investigating stress and song in song sparrows have shown that treatment with corticosterone, the hormone that mediates the stress response in birds, or food restriction during development did not lead to a difference in trill rate or song-type stereotypy but did lead to smaller syllable and song repertoires [56]. Further, adult assays of stress reactivity found a negative correlation between the magnitude of the acute stress response and song complexity as a combined measure of syllable and song repertoires [31]. Together, these findings suggest that, in song sparrows, metrics of song complexity, such as song repertoire, may be both more biologically meaningful and more responsive to early-life stress than metrics of song performance, like the rate of syllable production.

The high baseline level of complexity of song sparrow songs may prevent us from drawing solid conclusions about biological causes and consequences of any shifts in song using only mean song feature values across recordings, since, for example, dramatic changes in syntax or syllable composition could occur but not be reflected in the average values we analyzed. However, the observed shifts in certain features may hint at more nuanced changes happening in song or syllable structure. Future analysis of individual syllable types could probe these facets of song in greater depth.

There are several ways in which this study system and site may not be ideal for assessing song changes on a population scale. As with all community-science-based analyses, we lack metadata that would be useful in interpreting our analyses, such as the age of the bird in the recording. One potential challenge of studying severe environmental stress is that members of a population might conserve resources by reducing their reproductive output, for example by abandoning nest-building activities or failing to provision chicks. In the context of this study, this would mean that there would be very few juveniles developing during the height of the drought, and thus a minimal effect on the represented songs in the following years. However, the timing of the drought might have mitigated this challenge; since the drought did not reach its highest level of severity until mid-summer, many species would likely have already bred and raised chicks to at least fledgling age, increasing the likelihood that enough juveniles were exposed to drought stress to significantly affect population song.

To detect shifts in song that could potentially be linked to impaired learning due to the drought, as opposed to occurring due to normal stochasticity in population song, we compared the drought region to a control region; however, the best-sampled control regions we could find still had much smaller numbers of recordings than the drought region, reducing our power to detect any potential shifts in the songs in these control regions. Further, the only reasonably well-sampled potential control regions also experienced abnormally dry and moderate drought conditions, respectively, during 2016, since this drought was a regional phenomenon that affected much of the northeast United States.

Song in a population could shift significantly after an environmental stressor for reasons other than impaired learning. First, if there was heightened mortality in the population, songs could undergo a bottleneck effect akin to the stochastic loss of genetic diversity after extreme population reduction. In such a case, we might expect to see a narrower distribution of one or more of the song features after the reduction in population diversity that occurs during this type of bottleneck event. We did not observe such a shift in variance in any song features. Alternatively, if song features were correlated with another trait that was under selection pressure during drought conditions, for example body size, there could be indirect effects on song independent of learning ability. Second, a drought could induce individuals to emigrate from the region in pursuit of more hospitable conditions, or to not return to it after migration. If these individuals did not return in the following year once drought conditions had abated, others may disperse into those territories, changing the region’s song distribution by changing the members of the region. Third, adult songs, despite being crystallized and thus not expected to radically change, could theoretically be different during drought conditions, for example if the ambient temperature and humidity altered airflow through the syrinx or sound transmission through the air [57]. In this case, juveniles perceiving those song differences could hypothetically learn the song as perceived through that “filter” and propagate it to future pupils.

Other systems may prove more robust for testing the hypotheses and approaches we propose here. An ideal study system would likely be a long-term field study on a isolated or island population where historical data exist on rates of mortality and recruitment per year, especially during years when a stressful environmental event occurs, ample historical recordings of songs from years prior to the environmental stressor, and environmental data including water availability and a survey of available nutrient sources. If the species predominantly exhibits high-fidelity vertical learning, or learning from parent to offspring with little improvisation on the part of the pupil, having data on parentage and identity-linked song recordings would allow for comparisons between songs of the tutor and pupil directly. By measuring the similarity between these songs, it would be possible to more directly attribute any changes in song to mistakes in learning and track the frequency of mistakes over stressful and non-stressful years.

Studying a learned behavior, such as song, in the context of population-wide evolutionary pressures is particularly interesting since the resulting cultural evolutionary dynamics could be significantly more rapid than genetic evolution. Even without selection for a novel song type, the combination of drift and oblique transmission—from an adult to an unrelated juvenile—could lead to fixation of novel song variants far more rapidly than possible with a genetic mutation. Whether or not transient learning deficits in a subset of a population could induce a persistent shift in song, and therefore influence the long-term trajectory of the population, likely would depend on several factors.

Here, we present a study of the songs of two well-recorded species in a region that experienced a severe drought compared to a region that did not, and we observe a suite of song changes before versus after the time of the drought in only the species with the more complex song. If the trends we observed in the three years since the drought were to persist and a population’s song underwent a lasting change, there could be significant implications for conservation, particularly in the face of climate change that is likely to make severe weather events more common. If these changes were particularly salient in a given species’ song—for example, if the song changed in features that are important for mate choice—a population’s song could theoretically become unattractive to a sister population that maintained the typical ancestral song. This would suggest a potential side effect of anthropogenic climate change that has not been studied: that the sublethal experiences of individuals could impact the development of behaviors that are culturally transmitted to future generations, leading to an increase in reproductive isolation by sexual selection of this trait that is both plastic and sensitive to hostile conditions [58].

## References

1. Price TD, Grant PR, Gibbs HL, Boag PT. Recurrent patterns of natural selection in a population of Darwin’s finches. Nature. 1984;309: 787–789.

2. Donihue CM, Kowaleski AM, Losos JB, Algar AC, Baeckens S, Buchkowski RW, et al. Hurricane effects on Neotropical lizards span geographic and phylogenetic scales. Proc Natl Acad Sci U S A. 2020;117: 10429–10434.

3. Grant BR, Grant PR. Evolution of Darwin’s finches caused by a rare climatic event. Proceedings of the Royal Society of London Series B: Biological Sciences. 1993;251: 111–117.

4. Grant PR, Grant BR. Causes of lifetime fitness of Darwin’s finches in a fluctuating environment. Proc Natl Acad Sci U S A. 2011;108: 674–679.

5. Brown CR, Brown MB. INTENSE NATURAL SELECTION ON BODY SIZE AND WING AND TAIL ASYMMETRY IN CLIFF SWALLOWS DURING SEVERE WEATHER. Evolution. 1998;52: 1461–1475.

6. Sepers B, van den Heuvel K, Lindner M, Viitaniemi H, Husby A, van Oers K. Avian ecological epigenetics: pitfalls and promises. J Ornithol. 2019;160: 1183–1203.

7. Henry LP, Bruijning M, Forsberg SKG, Ayroles JF. The microbiome extends host evolutionary potential. Nat Commun. 2021;12: 5141.

8. Caspi T, Johnson JR, Lambert MR, Schell CJ, Sih A. Behavioral plasticity can facilitate evolution in urban environments. Trends Ecol Evol. 2022;37: 1092–1103.

9. Boyce WT, Sokolowski MB, Robinson GE. Genes and environments, development and time. Proc Natl Acad Sci U S A. 2020;117: 23235–23241.

10. Brass DP, Cobbold CA, Ewing DA, Purse BV, Callaghan A, White SM. Phenotypic plasticity as a cause and consequence of population dynamics. Ecol Lett. 2021;24: 2406–2417.

11. Gienapp P, Laine VN, Mateman AC, van Oers K, Visser ME. Environment-Dependent Genotype-Phenotype Associations in Avian Breeding Time. Front Genet. 2017;8: 102.

12. Regan CE, Beck KB, McMahon K, Crofts S, Firth JA, Sheldon BC. Social phenotype-dependent selection of social environment in wild great and blue tits: an experimental study. Proc Biol Sci. 2022;289: 20221602.

13. Marler P, Isaac D. Physical Analysis of a Simple Bird Song as Exemplified by the Chipping Sparrow. Condor. 1960;62: 124–135.

14. Kroodsma DE, Parker LD. Vocal Virtuosity in the Brown Thrasher. Auk. 1977;94: 783–785.

15. Robinson CM, Snyder KT, Creanza N. Correlated evolution between repertoire size and song plasticity predicts that sexual selection on song promotes open-ended learning. Elife. 2019;8. doi:10.7554/eLife.44454

16. Creanza N, Fogarty L, Feldman MW. Cultural niche construction of repertoire size and learning strategies in songbirds. Evol Ecol. 2016;30: 285–305.

17. Catchpole CK, Dittami J, Leisler B. Differential responses to male song repertoires in female songbirds implanted with oestradiol. Nature. 1984;312: 563–564.

18. Searcy WA. Song repertoire size and female preferences in song sparrows. Behav Ecol Sociobiol. 1984;14: 281–286.

19. Reid JM, Arcese P, Cassidy ALEV, Hiebert SM, Smith JNM, Stoddard PK, et al. Song repertoire size predicts initial mating success in male song sparrows, Melospiza melodia. Anim Behav. 2004;68: 1055–1063.

20. Nowicki S, Searcy WA. Song function and the evolution of female preferences: why birds sing, why brains matter. Ann N Y Acad Sci. 2004;1016: 704–723.

21. Brumm H, Zollinger SA, Slater PJB. Developmental stress affects song learning but not song complexity and vocal amplitude in zebra finches. Behav Ecol Sociobiol. 2009;63: 1387–1395.

22. MacDougall-Shackleton SA, Spencer KA. Developmental stress and birdsong: current evidence and future directions. J Ornithol. 2012;153: 105–117.

23. Svoboda M, LeComte D, Hayes M, Heim R, Gleason K, Angel J, et al. THE DROUGHT MONITOR. Bull Am Meteorol Soc. 2002;83: 1181–1190.

24. Sweet SK, Wolfe DW, DeGaetano A, Benner R. Anatomy of the 2016 drought in the Northeastern United States: Implications for agriculture and water resources in humid climates. Agric For Meteorol. 2017;247: 571–581.

25. Wingfield JC, Sullivan K, Jaxion-Harm J, Meddle SL. The presence of water influences reproductive function in the song sparrow (Melospiza melodia morphna). Gen Comp Endocrinol. 2012;178: 485–493.

26. Prior NH, Heimovics SA, Soma KK. Effects of water restriction on reproductive physiology and affiliative behavior in an opportunistically-breeding and monogamous songbird, the zebra finch. Horm Behav. 2013;63: 462–474.

27. Kozlovsky DY, Branch CL, Pitera AM, Pravosudov VV. Fluctuations in annual climatic extremes are associated with reproductive variation in resident mountain chickadees. R Soc Open Sci. 2018;5: 171604.

28. Langin KM, Sillett TS, Yoon J, Sofaer HR, Morrison SA, Ghalambor CK. Reproductive consequences of an extreme drought for orange-crowned warblers on Santa Catalina and Santa Cruz islands. Proceedings of the Seventh California Islands Symposium. Institute for Wildlife Studies; 2009.

29. Boag PT, Grant PR. The classical case of character release: Darwin’s finches (Geospiza) on Isla Daphne Major, Galápagos. Biol J Linn Soc Lond. 1984;22: 243–287.

30. Roberts LJ, Burnett R, Tietz J, Veloz S. Recent drought and tree mortality effects on the avian community in southern Sierra Nevada: a glimpse of the future? Ecol Appl. 2019;29: e01848.

31. Schmidt KL, Furlonger AA, Lapierre JM, MacDougall-Shackleton EA, MacDougall-Shackleton SA. Regulation of the HPA axis is related to song complexity and measures of phenotypic quality in song sparrows. Horm Behav. 2012;61: 652–659.

32. Hughes M, Nowicki S, Searcy WA, Peters S. Song-type sharing in song sparrows: implications for repertoire function and song learning. Behav Ecol Sociobiol. 1998;42: 437–446.

33. Ferree ED. Geographic variation in morphology of Dark-eyed Juncos and implications for population divergence. Wilson J Ornithol. 2013;125: 454–470.

34. Searfoss AM, Pino JC, Creanza N. Chipper: Open-source software for semi-automated segmentation and analysis of birdsong and other natural sounds. Methods Ecol Evol. 2020;11: 524–531.

35. Brenowitz EA, Beecher MD. Song learning in birds: diversity and plasticity, opportunities and challenges. Trends Neurosci. 2005;28: 127–132.

36. Weatherhead PJ, Forbes MRL. Natal philopatry in passerine birds: genetic or ecological influences? Behav Ecol. 1994;5: 426–433.

37. Liebgold EB, Gerlach NM, Ketterson ED. Similarity in temporal variation in sex-biased dispersal over short and long distances in the dark-eyed junco, Junco hyemalis. Mol Ecol. 2013;22: 5548–5560.

38. Huberty AF, Denno RF. PLANT WATER STRESS AND ITS CONSEQUENCES FOR HERBIVOROUS INSECTS: A NEW SYNTHESIS. Ecology. 2004. pp. 1383–1398. doi:10.1890/03-0352

39. White TCR. The abundance of invertebrate herbivores in relation to the availability of nitrogen in stressed food plants. Oecologia. 1984;63: 90–105.

40. Del Hoyo J, Del Hoyo J, Elliott A, Sargatal J. Handbook of the birds of the world. Lynx edicions Barcelona; 1992.

41. Cardoso GC, Atwell JW, Ketterson ED, Price TD. Song types, song performance, and the use of repertoires in dark-eyed juncos (Junco hyemalis). Behav Ecol. 2009;20: 901–907.

42. Williams L, MacRoberts MH. Individual Variation in Songs of Dark-Eyed Juncos. Condor. 1977;79: 106–112.

43. Newman MM, Yeh PJ, Price TD. Song variation in a recently founded population of the dark-eyed junco (junco hyemalis). Ethology. 2008;114: 164–173.

44. Wood WE, Yezerinac SM. Song Sparrow (Melospiza Melodia) Song Varies with Urban Noise. Auk. 2006;123: 650–659.

45. Marler P, Peters S. A sensitive period for song acquisition in the song sparrow, Melospiza melodia: A case of age-limited learning. Ethology. 1987;76: 89–100.

46. Borror DJ. Song Variation in Maine Song Sparrows. Wilson Bull. 1965;77: 5–37.

47. Podos J. A PERFORMANCE CONSTRAINT ON THE EVOLUTION OF TRILLED VOCALIZATIONS IN A SONGBIRD FAMILY (PASSERIFORMES: EMBERIZIDAE). Evolution. 1997;51: 537–551.

48. Ballentine B, Hyman J, Nowicki S. Vocal performance influences female response to male bird song: an experimental test. Behav Ecol. 2004;15: 163–168.

49. Cramer ERA. Vocal deviation and trill consistency do not affect male response to playback in house wrens. Behav Ecol. 2012;24: 412–420.

50. Caro SP, Sewall KB, Salvante KG, Sockman KW. Female Lincoln’s sparrows modulate their behavior in response to variation in male song quality. Behav Ecol. 2010;21: 562–569.

51. Searcy WA, Peters S, Kipper S, Nowicki S. Female response to song reflects male developmental history in swamp sparrows. Behav Ecol Sociobiol. 2010;64: 1343–1349.

52. O’Loghlen AL, Beecher MD. Sexual preferences for mate song types in female song sparrows. Anim Behav. 1997;53: 835–841.

53. O’Lochlen AL, Beecher MD. Mate, neighbour and stranger songs: a female song sparrow perspective. Anim Behav. 1999;58: 13–20.

54. Hernandez AM, Pfaff JA, MacDougall-Shackleton EA, MacDougall-Shackleton SA. The development of geographic song preferences in female song SparrowsMelospiza melodia. Ethology. 2009;115: 513–521.

55. Hiebert SM, Stoddard PK, Arcese P. Repertoire size, territory acquisition and reproductive success in the song sparrow. Anim Behav. 1989;37: 266–273.

56. Schmidt KL, Moore SD, MacDougall-Shackleton EA, MacDougall-Shackleton SA. Early-life stress affects song complexity, song learning and volume of the brain nucleus RA in adult male song sparrows. Anim Behav. 2013;86: 25–35.

57. Pandit MM, Bridge ES, Ross JD. Environmental conditions lead to shifts in individual communication, which can cause cascading effects on soundscape composition. Ecol Evol. 2022;12: e9359.

58. Mendelson TC, Safran RJ. Speciation by sexual selection: 20 years of progress. Trends Ecol Evol. 2021;36: 1153–1163.

