## Supplemental Tables and Figures for "Cultural signatures of punctuated environmental change: measuring the effects of population-wide stress on song distribution in dark-eyed juncos and song sparrows"

| Number of unique recordings- dark-eyed junco | Drought Region | Control Region |
|---|---|---|
| 2006-June 2016 (Pre-drought) | 131 bouts from 46 recordings (45 ML, 1 XC) | 11 bouts from 8 recordings (8 ML) |
| July 2016-Dec 2016 (During) | 14 bouts from 5 recordings (5 ML) | 0 |
| 2017-2019 (Post-drought) | 199 bouts from 73 recordings (64 ML, 1 XC, 8 self) | 62 bouts from 48 recordings (47 ML, 1 XC) |

**Table S1A: Distribution of recordings of dark-eyed juncos in the drought and control regions.**

| Number of unique recordings- song sparrow | Drought Region | Control Region |
|---|---|---|
| 2006-June 2016 (Pre-drought) | 409 bouts from 37 recordings (34 ML, 3 XC) | 72 bouts from 16 recordings (1 ML, 15 XC) |
| July 2016-Dec 2016 (During) | 9 bouts from 4 recordings (4 ML) | 9 bouts from 3 recordings (2 ML, 1 XC) |
| 2017-2019 (Post-drought) | 745 bouts from 159 recordings (148 ML, 4 XC, 7 self) | 330 bouts from 83 recordings (74 ML, 9 XC) |

**Table S1B: Distribution of recordings of song sparrows in the drought and control regions.**

| Song Feature | Log transformed | Shapiro-Wilk Before, Drought Region | Shapiro-Wilk After, Drought Region | Shapiro-Wilk Before, Control Region | Shapiro-Wilk After, Control Region | p-value Before vs After, Drought Region | p-value Before vs After, Control Region | Holm-Bonferroni Threshold, Drought Region | Significant After Bonferroni, Drought Region | Holm-Bonferroni Threshold, Control Region | Significant After Bonferroni, Control Region | Brown-Forsythe test for unequal variance, Before vs After, Drought Region | Brown-Forsythe test for unequal variance, Before vs After, Control Region |
|---|---|---|---|---|---|---|---|---|---|---|---|---|---|
| average minimum frequency of syllables | No | 0.975 | 0.724 | 0.287 | 0.240 | 0.788 | 0.849 | 0.0250 | FALSE | 0.0250 | FALSE | 0.653 | 0.192 |
| overall frequency range | No | 0.537 | 0.879 | 0.879 | 0.448 | 0.603 | 0.198 | 0.0100 | FALSE | 0.0063 | FALSE | 0.165 | 0.213 |
| number of notes per syllable | Yes | 0.050 | 0.479 | 0.178 | 0.045 | 0.040 | 0.898 | 0.0033 | FALSE | 0.0500 | FALSE | 0.032 | 0.527 |
| number of unique syllables | Yes | 5.25E-09 | 3.37E-09 | 0.016 | 2.40E-06 | 0.534 | 0.754 | 0.0071 | FALSE | 0.0250 | FALSE | 0.269 | 0.846 |
| average maximum frequency of syllables | Yes | 0.062 | 0.779 | 0.295 | 0.176 | 0.879 | 0.727 | 0.0250 | FALSE | 0.0167 | FALSE | 0.360 | 0.863 |
| mean syllable duration | Yes | 0.051 | 0.016 | 0.986 | 0.064 | 0.890 | 0.204 | 0.0500 | FALSE | 0.0083 | FALSE | 0.842 | 0.969 |
| standard deviation of syllable duration | Yes | 0.014 | 0.889 | 0.829 | 0.099 | 0.348 | 0.194 | 0.0063 | FALSE | 0.0071 | FALSE | 0.072 | 0.644 |
| rate of syllable production | Yes | 0.075 | 0.004 | 0.961 | 0.021 | 0.207 | 0.146 | 0.0056 | FALSE | 0.0063 | FALSE | 0.599 | 0.414 |
| total number of syllables | Yes | 0.380 | 0.010 | 0.542 | 0.938 | 0.069 | 0.060 | 0.0042 | FALSE | 0.0056 | FALSE | 0.259 | 0.846 |
| mean syllable stereotypy | Yes | 0.053 | 9.86E-05 | 0.972 | 0.027 | 0.702 | 0.034 | 0.0083 | FALSE | 0.0050 | FALSE | 0.252 | 0.379 |
| degree of syllable repetition | No | 0.552 | 0.332 | 0.160 | 0.123 | 0.098 | 0.015 | 0.0045 | FALSE | 0.0045 | FALSE | 0.417 | 0.255 |
| total number of notes | Yes | 0.793 | 0.754 | 0.625 | 0.006 | 0.005 | 0.005 | 0.0029 | FALSE | 0.0042 | FALSE | 0.966 | 0.639 |
| bout duration | Yes | 0.579 | 0.087 | 0.684 | 0.006 | 0.015 | 0.004 | 0.0031 | FALSE | 0.0038 | FALSE | 0.908 | 0.153 |

**Table S2. Statistical analysis of dark-eyed junco songs.** When all Shapiro-Wilk tests were not significant, indicating normality, we compared song features using a t-test. Otherwise, we log-transformed and used a Wilcoxon rank-sum test. We corrected for multiple hypothesis testing with a Holm-Bonferroni correction. No song features were significantly different before vs. after the drought in either the drought or control region.

| Song Feature | Log transformed | Shapiro-Wilk Before, Drought Region | Shapiro-Wilk After, Drought Region | Shapiro-Wilk Before, Control Region | Shapiro-Wilk After, Control Region | p-value Before vs After, Drought Region | p-value Before vs After, Control Region | Holm-Bonferroni Threshold, Drought Region | Significant After Bonferroni, Drought Region | Holm-Bonferroni Threshold, Control Region | Significant After Bonferroni, Control Region | Brown-Forsythe test for unequal variance, Before vs After, Drought Region | Brown-Forsythe test for unequal variance, Before vs After, Control Region |
|---|---|---|---|---|---|---|---|---|---|---|---|---|---|
| degree of syllable repetition | Yes | 0.452 | 0.273 | 0.093 | 0.447 | 5.00E-05 | 0.059 | 0.0036 | TRUE | 0.0036 | FALSE | 0.107 | 0.062 |
| number of notes per syllable | Yes | 0.491 | 3.13E-05 | 0.278 | 0.048 | 0.0006 | 0.267 | 0.0038 | TRUE | 0.0050 | FALSE | 0.753 | 0.481 |
| total number of syllables | No | 0.695 | 0.105 | 0.505 | 0.501 | 0.0026 | 0.020 | 0.0042 | TRUE | 0.0033 | FALSE | 0.551 | 0.359 |
| rate of syllable production | Yes | 0.043 | 0.571 | 0.860 | 0.282 | 0.0074 | 0.061 | 0.0045 | FALSE | 0.0038 | FALSE | 0.038 | 0.619 |
| mean syllable duration | Yes | 0.989 | 0.861 | 0.717 | 0.233 | 0.0199 | 0.089 | 0.0050 | FALSE | 0.0045 | FALSE | 0.086 | 0.284 |
| average minimum frequency of syllables | Yes | 0.510 | 0.017 | 0.238 | 0.534 | 0.1030 | 0.341 | 0.0056 | FALSE | 0.0071 | FALSE | 0.098 | 0.537 |
| standard deviation of syllable duration | Yes | 0.985 | 0.093 | 0.307 | 0.102 | 0.1788 | 0.270 | 0.0063 | FALSE | 0.0056 | FALSE | 0.048 | 0.859 |
| bout duration | Yes | 0.012 | 4.73E-04 | 0.240 | 0.279 | 0.2164 | 0.406 | 0.0071 | FALSE | 0.0125 | FALSE | 0.325 | 0.335 |
| duration of the longest syllable | Yes | 0.026 | 7.36E-05 | 0.306 | 0.013 | 0.2402 | 0.372 | 0.0083 | FALSE | 0.0083 | FALSE | 0.096 | 0.848 |
| total number of notes | Yes | 0.207 | 0.321 | 0.643 | 0.255 | 0.3965 | 0.808 | 0.0100 | FALSE | 0.0500 | FALSE | 0.182 | 0.263 |
| overall frequency range | Yes | 2.61E-04 | 1.49E-06 | 0.384 | 0.001 | 0.5581 | 0.328 | 0.0125 | FALSE | 0.0063 | FALSE | 0.954 | 0.184 |
| duration of the shortest syllable | Yes | 0.267 | 0.683 | 0.289 | 0.592 | 0.5730 | 0.087 | 0.0167 | FALSE | 0.0042 | FALSE | 0.409 | 0.492 |
| average maximum frequency of syllables | Yes | 0.188 | 0.005 | 0.340 | 0.068 | 0.8469 | 0.763 | 0.0250 | FALSE | 0.0250 | FALSE | 0.522 | 0.661 |
| number of unique syllables | Yes | 0.031 | 0.638 | 0.884 | 0.110 | 0.9474 | 0.381 | 0.0500 | FALSE | 0.0100 | FALSE | 0.634 | 0.361 |

**Table S3. Statistical analysis of song sparrow songs.** When all Shapiro-Wilk tests were not significant, indicating normality, we compared song features using a t-test. Otherwise, we log-transformed and used a Wilcoxon rank-sum test. We corrected for multiple hypothesis testing with a Holm-Bonferroni correction. Three song features were significantly different before vs. after the drought in the drought region, and no song features differed in the control region.

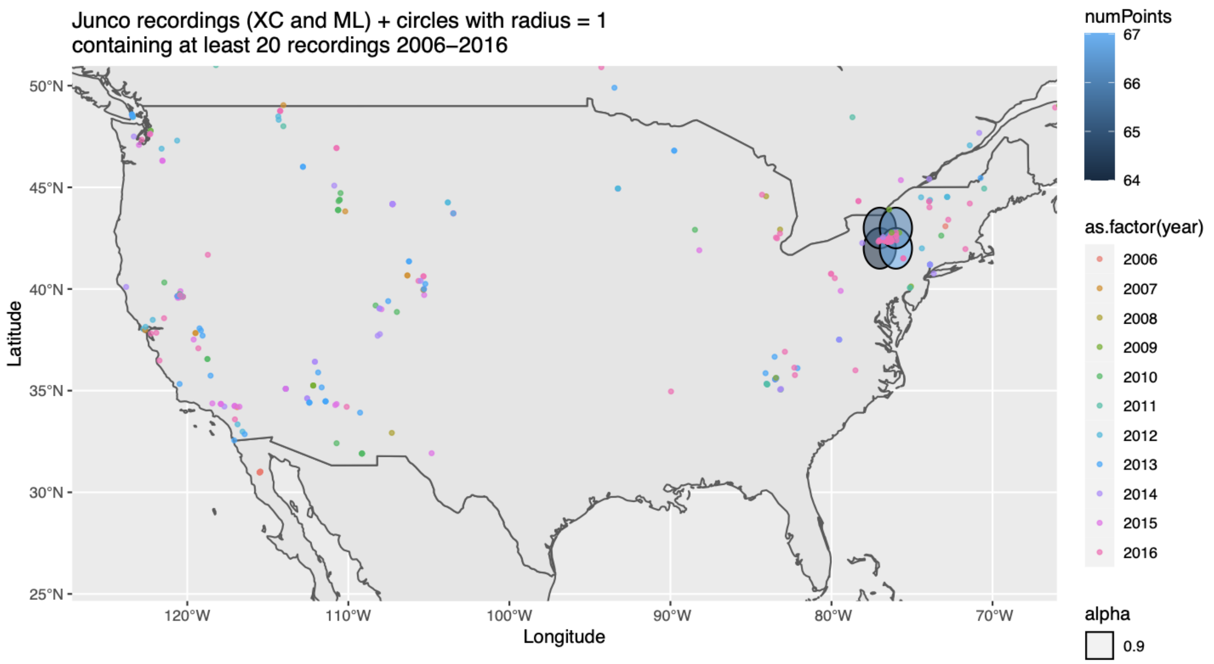


**Figure S1A: Dark-eyed junco recordings publicly available on Macaulay Library or xeno-canto recorded between 2006 and 2016.** Of all regions centered on each value of (Longitude, Latitude) ranging from 70W through 89W and 30N through 47N, the plotted, filled circle regions are the only regions that contain at least 20 recordings. The four circles shown here all include part of Tompkins County, NY.

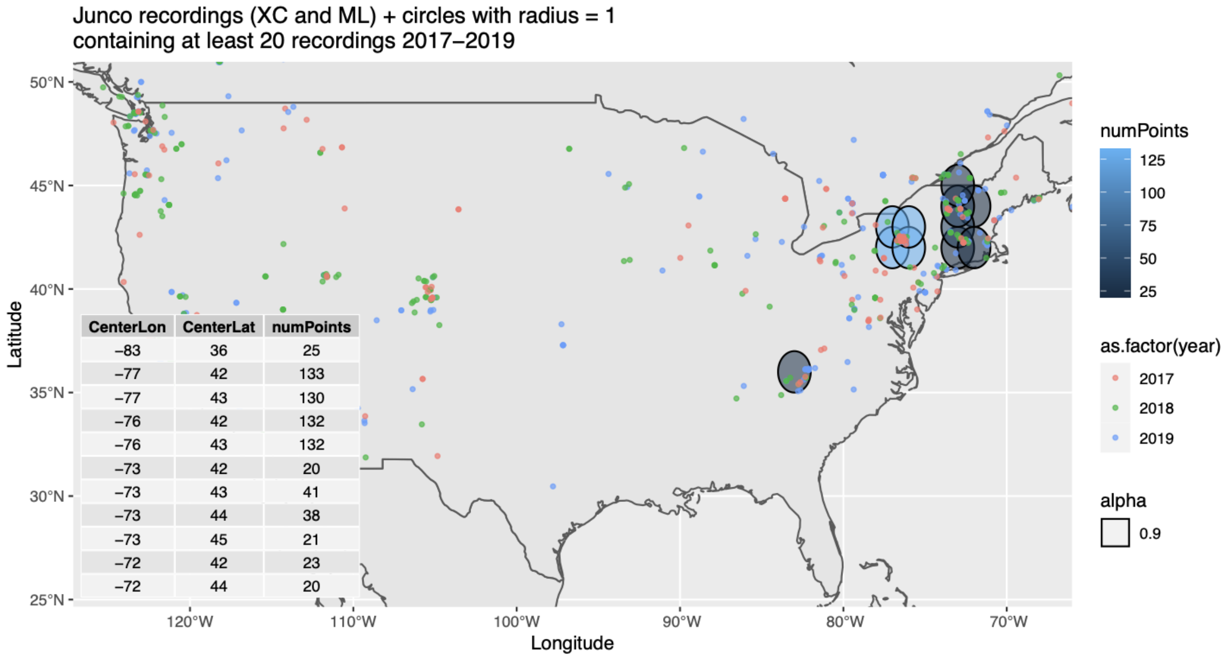


| CenterLon | CenterLat | numPoints |
|---|---|---|
| −83 | 36 | 25 |
| −77 | 42 | 133 |
| −77 | 43 | 130 |
| −76 | 42 | 132 |
| −76 | 43 | 132 |
| −73 | 42 | 20 |
| −73 | 43 | 41 |
| −73 | 44 | 38 |
| −73 | 45 | 21 |
| −72 | 42 | 23 |
| −72 | 44 | 20 |

**Figure S1B: Dark-eyed junco recordings publicly available on Macaulay Library or xeno-canto recorded between 2017 and 2019.** Of all regions centered on each value of (Longitude, Latitude) ranging from 70W through 89W and 30N through 47N, the plotted, filled circle regions are the only regions that contain at least 20 recordings. Inlaid table shows the exact counts for each circle and the (Longitude, Latitude) center of the plotted circle.

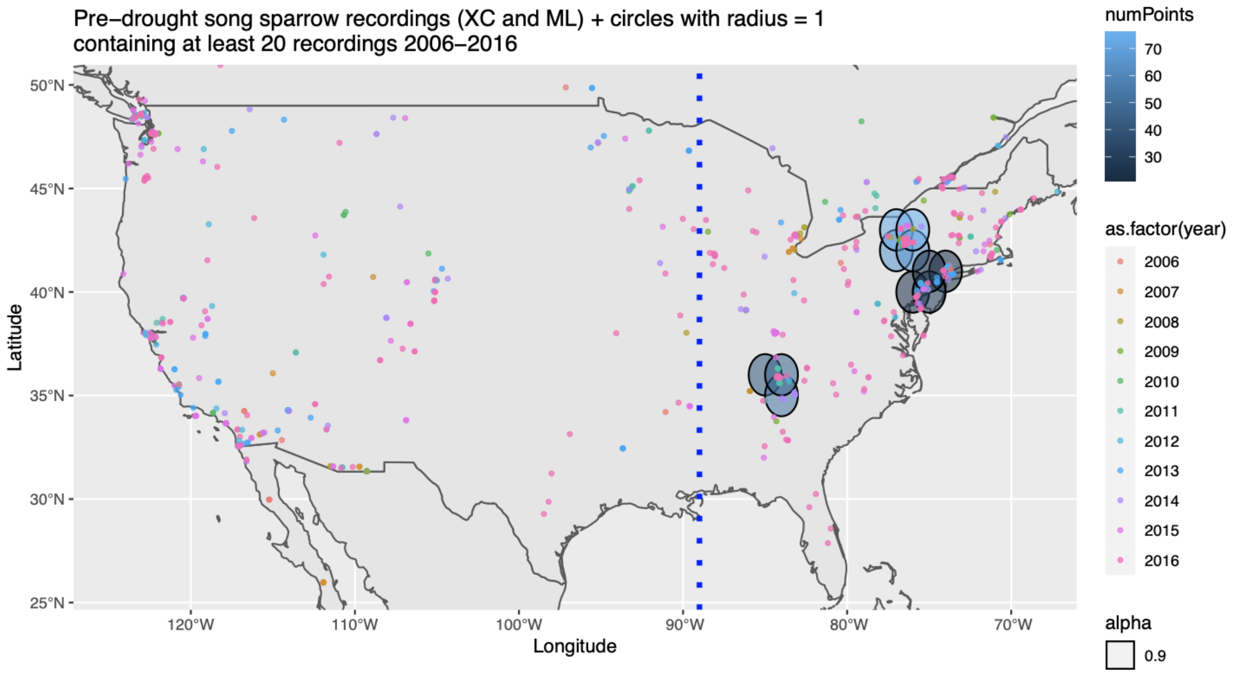


**Figure S1C: Song sparrow recordings publicly available on Macaulay Library or xeno-canto recorded between 2006 and 2016.** Of all regions centered on each value of (Longitude, Latitude) ranging from 70W through 89W and 30N through 47N, the plotted, filled circle regions are the only regions that contain at least 20 recordings.

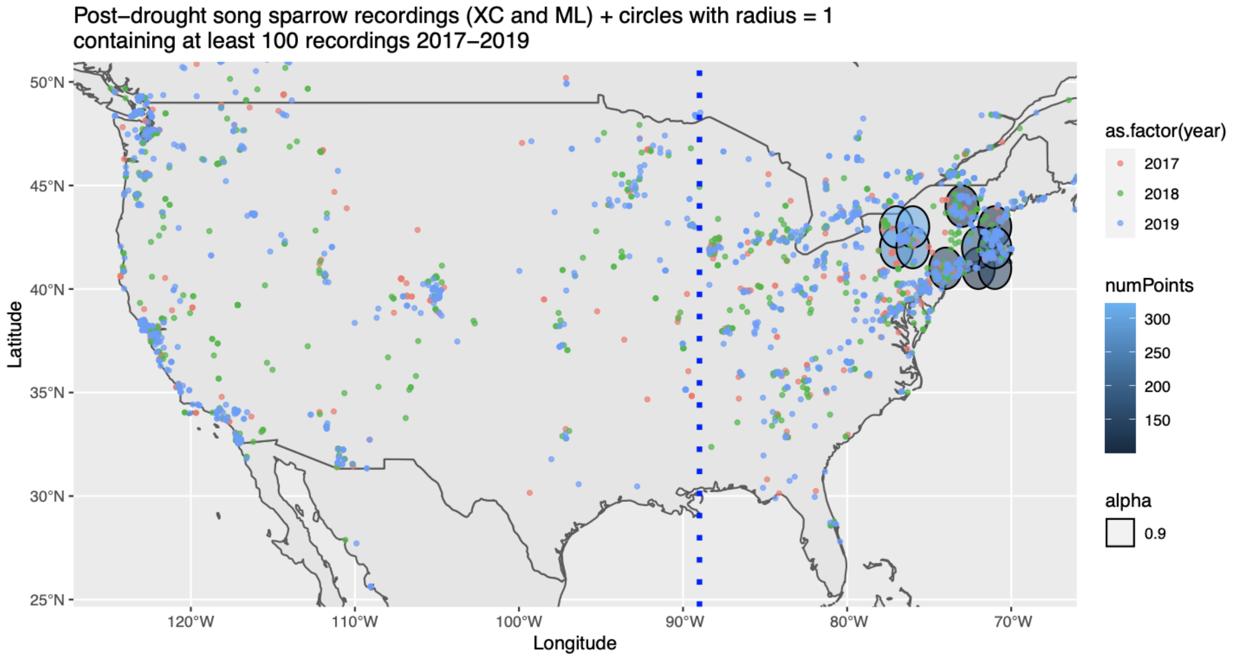


**Figure S1D: Song sparrow recordings publicly available on Macaulay Library or xeno-canto recorded between 2017 and 2019.** Of all regions centered on each value of (Longitude, Latitude) ranging from 70W through 89W and 30N through 47N, the plotted, filled circle regions are the regions that contain at least 100 recordings.

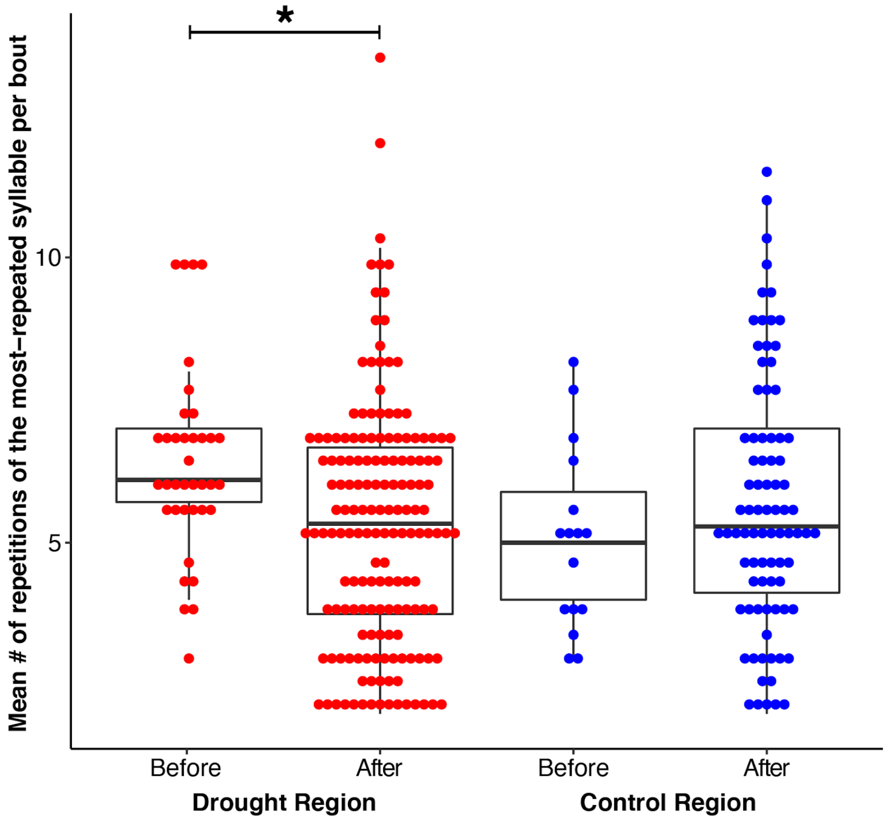


**Figure S2: Distributions of average counts of the most-repeated syllable per bout per recording within song sparrow populations before and after the 2016 drought**. For each bout, we found the most repeated syllable type based on the syllable type assignment from the Chipper analysis, counted the number of times the most repeated syllable was produced, and averaged those counts across all bouts sampled from a given recording. Overall, in recordings after the drought, the most-repeated syllable type per bout was repeated fewer times on average (t-test $p = 0.007$).
